# The interface of condensates of the hnRNPA1 low complexity domain promotes formation of amyloid fibrils

**DOI:** 10.1101/2022.05.23.493075

**Authors:** Miriam Linsenmeier, Lenka Faltova, Umberto Capasso Palmiero, Charlotte Seiffert, Andreas M. Küffner, Dorothea Pinotsi, Jiangtao Zhou, Raffaele Mezzenga, Paolo Arosio

## Abstract

The maturation of liquid-like protein condensates into amyloid fibrils has been associated with neurodegenerative diseases. Here, we analyze the amyloid formation mediated by condensation of the low-complexity domain of hnRNPA1, a protein involved in Amyotrophic Lateral Sclerosis (ALS). We show that phase separation and fibrillation are connected but distinct processes which are independently mediated by different regions of the protein sequence. By monitoring the spatial and temporal evolution of amyloid formation we demonstrate that the formation of fibrils does not occur homogeneously inside the droplets but is promoted at the interface of the condensates. Consistently, we further show that coating the interface of the droplets with surfactant molecules inhibits fibril formation. Our results indicate that the interface of biomolecular condensates can act as an important catalyst for fibril formation, and therefore could represent a possible therapeutic target against the formation of aberrant amyloids mediated by condensation.

## Introduction

Recent findings indicate that cells can regulate functions in space and time *via* membraneless organelles composed of proteins and nucleic acids.^1–3^ These viscoelastic condensates^4^ exhibit a variety of material properties, manifesting physical states that range from liquid-like^5–7^ to dynamically arrested^8,9^.

In some cases dynamic condensates exhibit a maturation process over time towards either arrested states^8,10,11^ or amyloid fibrils^12–15^. While for some condensates this “aging” can be functional^16^, in other cases liquid to solid transitions can lead to aberrant behavior, as often observed with maturation into amyloids^17^.

Transitions from liquid-like condensates to amyloids have been observed for peptides^18,19^, α -synuclein^20^, tau^21,22^ and a variety of RNA-binding proteins (RBPs) associated with Amyotrophic Lateral Sclerosis (ALS) and Frontotemporal Dementia (FTD)^23,24^, including Transactive response DNA binding protein 43 (TDP-43)^25^, Fused in sarcoma (FUS)^12^, and heterogeneous nuclear ribonucleoprotein A1/2 (hnRNPA1/2)^13^.

These RBPs have been found in insoluble cytoplasmic inclusions in neurons of *post-mortem* brain and spinal cord tissue of ALS patients^23,26–28^ and have been shown to form amyloid fibrils with characteristic cross-β-sheet architecture^29^ *in vitro*^30–32^ and *ex vivo*^33^. These proteins have also been associated with the formation of several membraneless organelles including stress granules and paraspeckles^13,34^.

Recent effort has been focusing on unraveling the molecular modulators of the liquid-amyloid transitions of different RBPs. In this context, *in vitro* studies have shown that ALS-associated amino acid substitutions and post-translational modifications affect the liquid-amyloid transition of these RBPs. For instance, the balance of FUS phase separation and amyloid formation is modulated by phosphorylation and acetylation^12,35,36^. These findings have supported the hypothesis that liquid-amyloid transitions could play an important role in some amyloid-associated diseases.

Despite the evidence that the formation of liquid-like condensates or the recruitment of aggregation-prone proteins therein could be an event promoting fibril formation^37,38^, the molecular mechanisms underlying such liquid-amyloid transitions are only starting to be understood. Unraveling these mechanisms has huge potential to identify effective therapeutic strategies to prevent the formation of aberrant fibrils^39^.

In this work, we analyze the liquid-amyloid transition of the intrinsically disordered region of hnRNPA1. hnRNPA1 consists of a globular domain containing two RNA-recognition motifs (RRM1 and RRM2) and a C-terminal intrinsically disordered region, indicated as low-complexity domain (LCD). Both the liquid-liquid phase separation (LLPS) and the formation of amyloid fibrils of hnRNPA1 have been shown to be largely governed by the LCD^13,23^. Moreover, single point mutations, frameshifts and extensions of the LCD have been identified in ALS patients and associated not only with mis-localization of the protein from the nucleus to the cytoplasm and subsequent accumulation therein, but also with altered LLPS, fibrillization and stress granule dynamics^23,40^.

The sequence of the LCD is very intriguing because it exhibits a peculiar “molecular grammar” and contains regions that drive LLPS and others that promote amyloid formation. Specifically, LLPS is mediated by regularly spaced, aromatic amino acids that act as “sticker” moieties^41,42^, while amyloid formation is mediated by LARKS (low-complexity aromatic-rich kinked segments)^43,44^ that promote β-sheet structures and reversible hydrogels at high protein concentrations *in vitro*^45^. This LCD is therefore an ideal model system to probe the correlation between LLPS and amyloid formation.

Here, we demonstrate that LLPS and amyloid formation of the LCD of hnRNPA1 are correlated but distinct processes mediated by different regions of the protein and that amyloid formation is largely accelerated when initiated from condensates. By monitoring the temporal and spatial evolution of fibrillation with a variety of biophysical techniques, we show that the formation of hnRNPA1 amyloid fibrils from condensates is promoted at the interface of the droplets. We further show that targeting the interface is an efficient way to inhibit this liquid-amyloid transition.

## Results

### The liquid-liquid phase separation and amyloid formation of the hnRNPA1-B LCD are independently mediated by different regions of the protein sequence

We investigate the C-terminal LCD of two splicing variants of hnRNPA1, isoforms hnRNPA1-A (A-LCD) and hnRNPA1-B (B-LCD), the latter containing 52 additional amino acids. The sequences of the proteins are shown in Fig. 1A, which highlights the residues that act as stickers and are responsible for LLPS^41^, the regions that are prone to form amyloids (see Materials and Methods and Suppl. Table 1).

**Figure 1:**
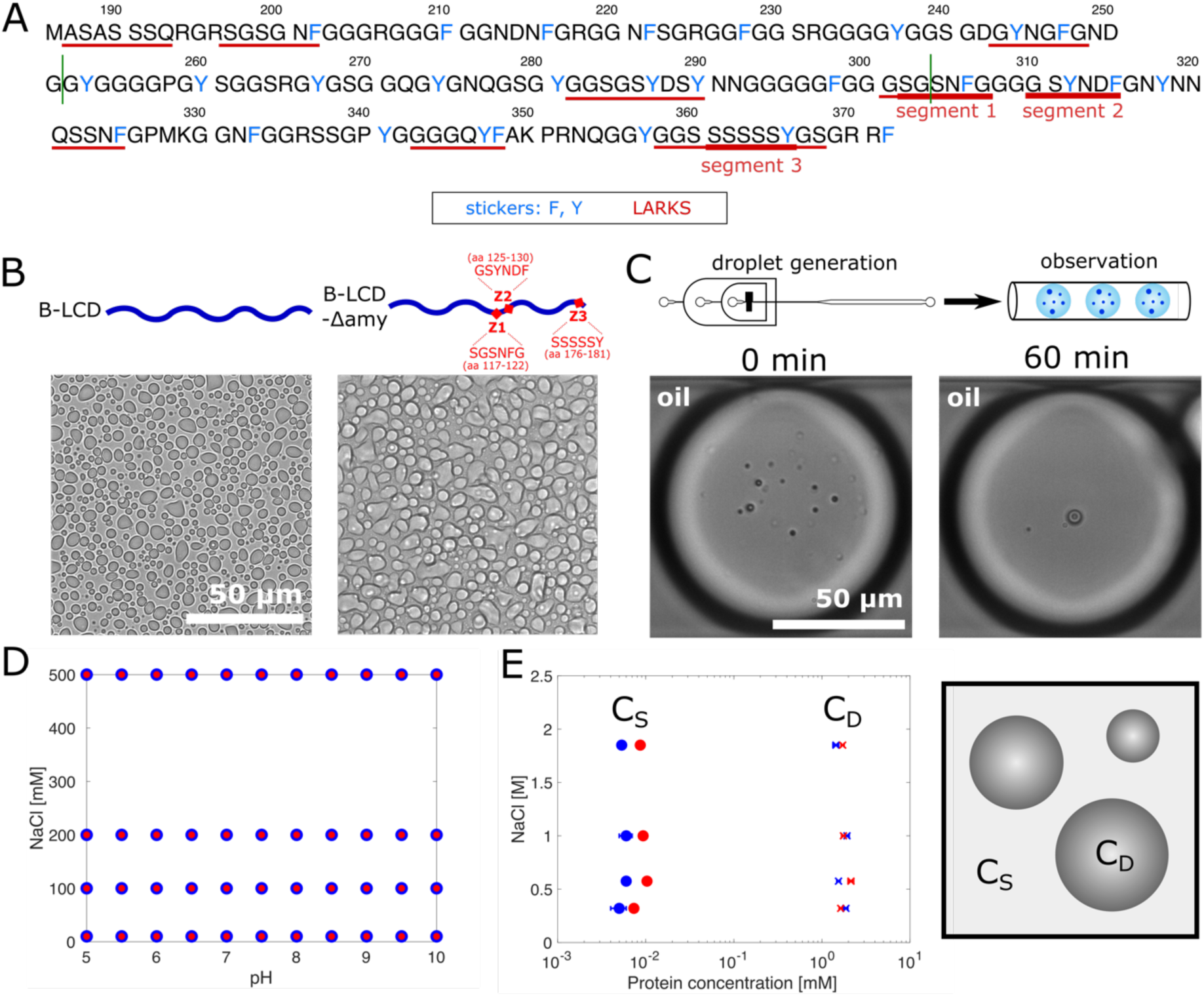
Liquid-liquid phase separation of B-LCD and zipper-depleted variant B-LCD-Δamy. **(A)** Sequence of the low-complexity domain of hnRNPA1-B (B-LCD). LCD of hnRNPA1-A (A-LCD) comprises amino acids 186-250 and 303-372 (borders indicated by green lines). Stickers (aromatic amino acids) are indicated in blue while fibril forming segments (steric zippers / LARKS) are underlined in red. **(B)** Brightfield microscopy images of droplets formed by B-LCD and zipper-depleted construct B-LCD-Δamy lacking three fibril forming segments (ΔSGSNFG, ΔGSYNDF, ΔSSSSSY). **(C)** B-LCD droplets are liquid-like and merge in microfluidic water-in-oil droplet compartments until one single condensate is formed within 60 min. **(D)** Both B-LCD (blue circles) and B-LCD-Δamy (red circles) undergo LLPS under a broad range of ionic strength and pH values. Circles indicate presence of LLPS. **(E)** Protein concentrations inside (C_D_) and outside (C_S_) the droplets at different ionic strengths are similar for B-LCD (blue) and B-LCD-Δamy (red). Error bars represent the standard deviation of the average protein concentration inside and outside the droplets, obtained by measuring 3 different droplets (for C_D_) and 3 independent (for C_S_) samples at each NaCl concentration. When not visible, the error bars are smaller than the symbol

In addition to the A- and B-LCD sequences, we generated a mutant B-LCD in which we eliminated three regions promoting amyloid formation: segment 1 (S1, ΔSGSNFG), which rapidly forms amyloid fibrils on its own^46^; segment 2 (S2, ΔGSYNDF) whose deletion in the LCD of isoform A effectively inhibits amyloid formation^13,47^; and segment 3 (S3, ΔSSSSSY), which exhibits the lowest interaction free energy according to ZipperDB (Suppl. Table 1). Segments S1 and S2 (Fig. 1A) overlap with the fibril core previously identified by cryo-electron microscopy^31^, and segments S1 and S3 have been predicted to exhibit LARK-like properties (LARKSdb^43^). In the following we indicate this variant depleted of three fibril-forming segments S1-S3 as B-LCD-Δamy.

All variants exhibited liquid-liquid phase separation over a broad range of buffer conditions, forming liquid-like droplets (Fig. 1B, Suppl. Fig. S1). We encapsulated the protein solution in water-in-oil microfluidic droplet compartments, which allows to accurately monitor coarsening events^48,49^. LCD droplets underwent coalescence until one single condensate per water-in-oil compartment was observed (Fig. 1C), therefore confirming the liquid-like nature of the condensates.

Importantly, the LLPS of all variants exhibited the same dependence on pH value and ionic strength (Fig. 1D). Additionally, we measured the concentration of the protein outside (C_S_) and inside (C_D_) the droplets at different ionic strengths. The soluble protein concentration C_S_ was measured by UV absorbance after separating the droplet phase by centrifugation. The C_D_ was measured by Raman spectroscopy, using a characteristic phenylalanine peak as standard^50^ (see Materials and Methods). We observed very similar values for B-LCD and B-LCD-Δamy (Fig. 1E), confirming that the phase separation of the two constructs is nearly identical.

After characterizing the LLPS of the LCD variants at time zero, we monitored the formation of fibrils over time by a variety of biophysical techniques, including Thioflavin T (ThT) staining, optical microscopy, re-scan confocal microscopy, atomic force microscopy and Raman spectroscopy (Fig. 2, Suppl. Fig. S2). For both the A-LCD and B-LCD we observed an increase of the fluorescence ThT signal over time which follows the characteristic sigmoidal profile observed during the kinetics of amyloid formation (Fig. 2A and Suppl. Fig. S3), reflecting the increase in β-sheet-rich structures over time. By contrast, the increase in ThT signal over time for B-LCD-Δamy was negligible (Fig. 2A).

**Figure 2:**
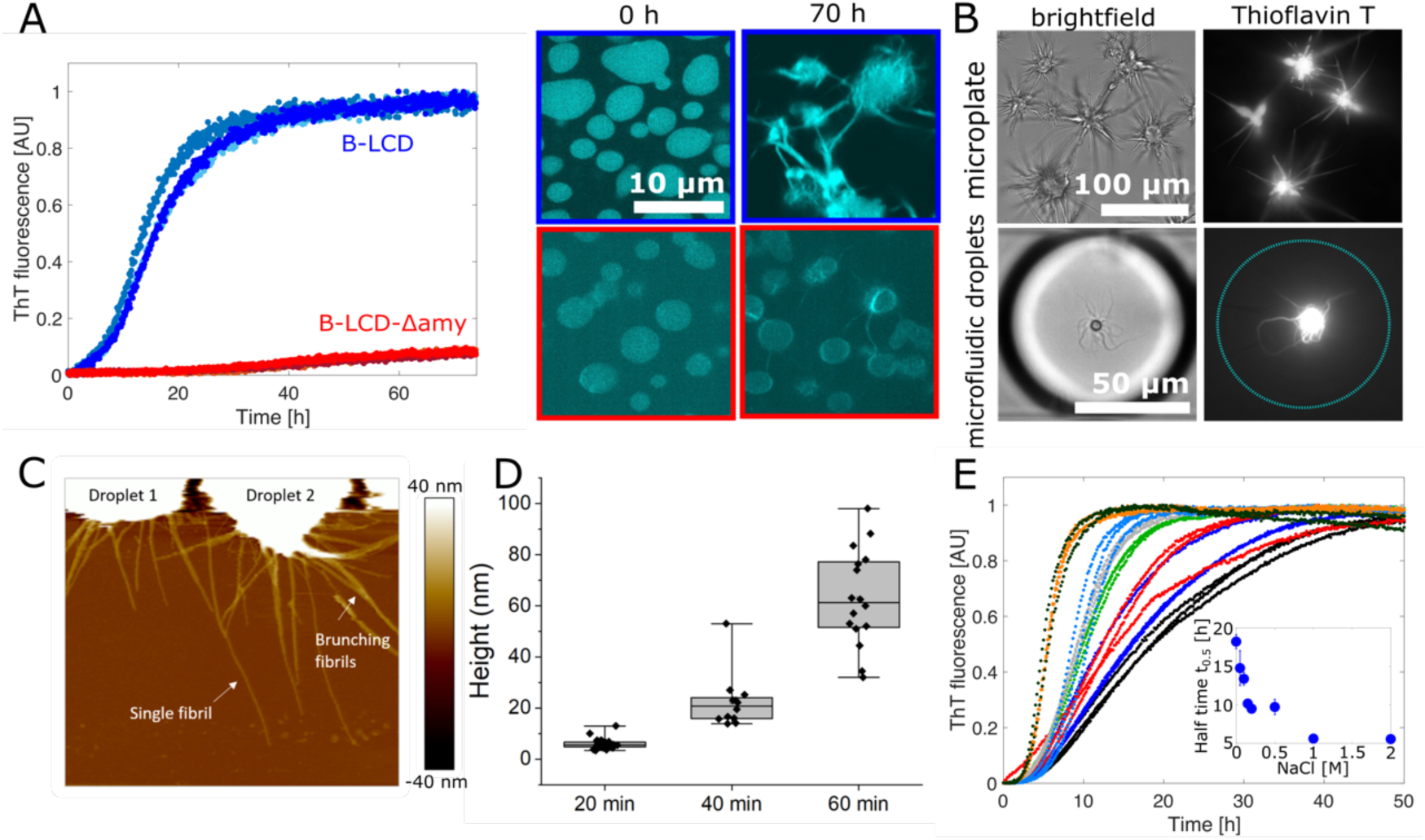
Liquid-amyloid transition of hnRNPA1-B is governed by fibril forming segments. **(A)** Fluorescence Thioflavin T (ThT) signal over time for B-LCD (blue) and B-LCD-Δamy (red). Re-scan confocal microscopy images at time zero and after 70 hours of incubation for the two constructs. Each curve represents the normalized ThT fluorescence of one independent sample. **(B)** ThT-stainable star-shaped aggregates formed by B-LCD in microplates and in microfluidic water-in-oil droplets imaged in brightfield and by widefield fluorescence microscopy after 6 days of incubation. **(C)** Atomic Force Microscopy (AFM) of star-shaped aggregates formed by B-LCD after 1 h. **(D)** Increase of fibril height over 60 min after formation, extracted from AFM images. **(E)** Acceleration of amyloid formation in 30 µM B-LCD sample with increasing NaCl concentration. Inset: half time t_0.5_ values as a function of NaCl concentration (0 mM NaCl – black, 50 mM NaCl – blue, 100 mM NaCl – red, 150 mM NaCl – light green, 200 mM NaCl – grey, 500 mM NaCl – light blue, 1 M NaCl – orange, 2 M NaCl – dark green).

These results were consistent with re-scan confocal microscopy images acquired at the end of the incubation, which show the presence of ‘starburst structures’^12^ in A-LCD and B-LCD samples, and considerably fewer fibrils for B-LCD-Δamy (Fig. 2A, Suppl. Fig. S3). The ‘starburst structures’ formed by A-LCD and B-LCD in both bulk assays and microfluidic droplets (Fig. 2B) are stainable by ThT and have been confirmed by AFM analysis (Fig. 2C). Further analysis of AFM images revealed that after 20 min the fibril height is approximately 6 nm, corresponding to the typical diameter of an individual amyloid fibril (Fig. 2D). Over time, fibrils grow into thicker aggregates (Fig. 2D).

We further monitored the changes in the secondary structure of the protein over time by Raman spectroscopy. For B-LCD, spectra acquired at time 0 (droplets) and after several days (star-shaped aggregates) showed a transition from a random coil state to β-sheet-rich structures^51,52^, while negligible changes were observed for B-LCD-Δamy (Suppl. Fig. S2).

Overall, the comparison between the LLPS behavior (Fig. 1) and the liquid-amyloid transition (Fig. 2) of B-LCD and B-LCD-Δamy indicates that the two processes are mediated by different regions of the LCD: the deletion of segments that strongly drive the formation of amyloid fibrils does not impact phase separation.

This result is further supported by aggregation profiles measured at increasing concentrations of salt. While the LLPS is largely independent of the ionic strength of the buffer (Fig. 1E), amyloid formation is accelerated by increasing salt concentrations (Fig. 2E), indicating that the interactions responsible for liquid-amyloid transition and LLPS are different and can be independently modulated. The increase in ionic strength likely neutralizes an electrostatic repulsive interaction mediated by the residue D314 (D262 in the A-LCD) in segment S2, therefore accelerating fibril formation. This mechanism is consistent with the mutation D314V, which accelerates amyloid formation by substitution of this charged residue^23^.

### Amyloid formation from liquid condensates of the hnRNPA1-B LCD is promoted at the interface

We next investigated how biomolecular condensation promotes the formation of amyloid fibrils. Since the kinetics of the liquid-amyloid transition of B-LCD was slower compared to A-LCD (Suppl. Fig. S3), in the following we decided to focus on B-LCD to better capture the transition process.

A first obvious effect of condensation is the local increase of the protein concentration, which from a thermodynamic point of view can exceed the critical concentration required to form amyloids^53^. Moreover, from a kinetic angle, the increase in concentration accelerates all microscopic nucleation and elongation reactions responsible for the formation of amyloids^53^.

We incubated B-LCD samples at concentrations below the LLPS critical concentration (∼ 2.5 µM, as shown in Fig. 3A) and we did not observe formation of amyloids over seven days of incubation (Fig. 3B), indicating that the formation of condensates is required to promote fibril formation (increasing the protein concentration over the critical concentration for liquid-solid transition) or at least strongly accelerates the process (kinetic effect). The absence of fibrils in samples containing B-LCD concentrations below the sub-critical concentrations for LLPS was further confirmed by transmission electron microscopy (TEM) analysis (Suppl. Fig. S4).

**Figure 3:**
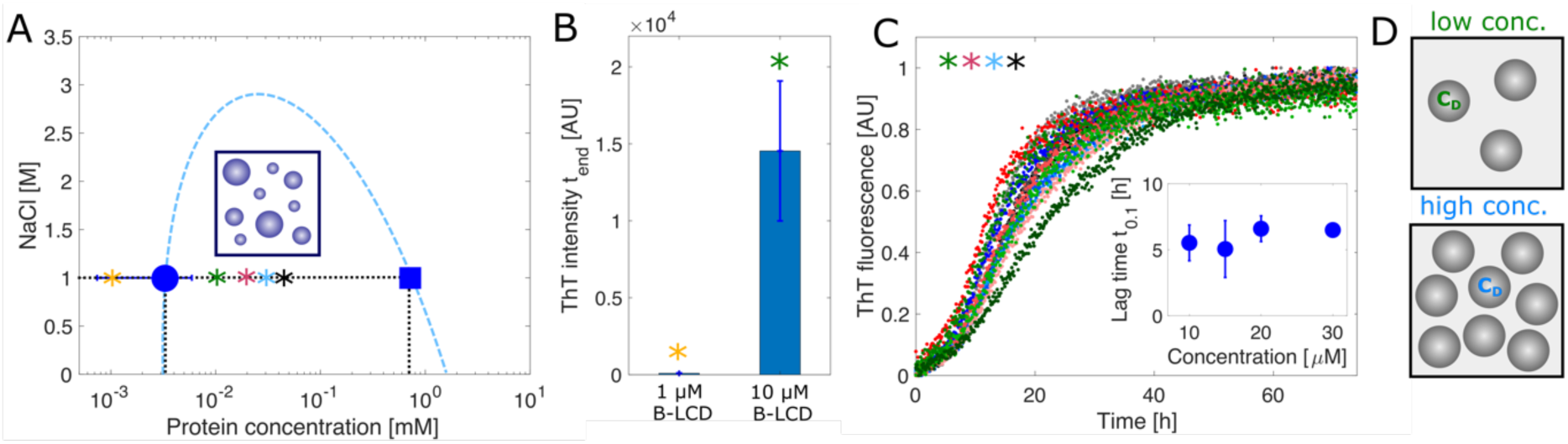
Condensation of hnRNPA1-B LCD accelerates amyloid formation. **(A)** Protein concentration inside (C_D_, blue square) and outside (C_S_, blue circle) the droplets. Asterisks indicate initial protein concentration in kinetic experiments in (B) and (C): 1 µM (yellow), 10 µM (green), 15 µM (red), 20 µM (blue), 30 µM (black). Error bars represent the standard deviation of the average protein concentration inside and outside the droplets, obtained by measuring 3 different droplets (for C_D_) and 3 independent samples (for C_S_) at each C_0_. The blue dashed line represents a guide to the eyes. **(B)** ThT fluorescence intensity value of 1 µM and 10 µM B-LCD solutions after 7 days incubation, respectively below and above the critical concentration of LLPS (C_c_ = 2.5 µM). **(C)** Normalized ThT kinetic profiles corresponding to the initial protein concentrations C_0_ in (A). The inset shows the average lag times t_0.1_ at various initial protein concentrations C_0_. Error bars represent standard deviation of three independent samples. **(D)** Droplets formed at low and high initial B-LCD concentration exhibit similar internal protein concentration (C_D_) but increase in number when the initial protein concentration is increased.

By contrast, samples that contained B-LCD at concentrations above the critical LLPS concentration showed fibril formation within 2 days (Fig. 3A-C). Normalized ThT profiles collected at different initial protein concentrations exhibited very similar kinetic profiles (Fig. 3C). In contrast to the formation of amyloids in bulk homogeneous solutions, these data show that the lag phase is independent of the initial protein concentration when amyloid formation is initiated from a condensate state. This result can be explained by the fact that the protein concentration inside the droplets C_D_ is constant and independent of the initial protein concentration (C_0_) (Fig. 3A). Each individual droplet behaves as an individual microreactor at constant concentration C_D_. Increasing the initial protein concentration leads to an increase in the number of droplets (Suppl. Fig. S5) (leading to an increase of the total mass of fibrils formed) but does not change the protein concentration inside the droplets, and therefore does not affect the lag phase. In the investigated concentration range, we did not observe any significant change of the size distribution of droplets at different initial protein concentrations C_0_ (Fig. 3D and Suppl. Fig. S5).

In addition to the temporal evolution of fibrillization, we next explored the spatial evolution of amyloid formation by acquiring microscopy images at different time points, while simultaneously monitoring the Thioflavin T profile (Fig. 4A). During the lag phase, microscopy images showed the formation of a ThT-positive rim on the droplet surface, which grew into star-shaped aggregates during the growth phase (Fig. 4A).

**Figure 4:**
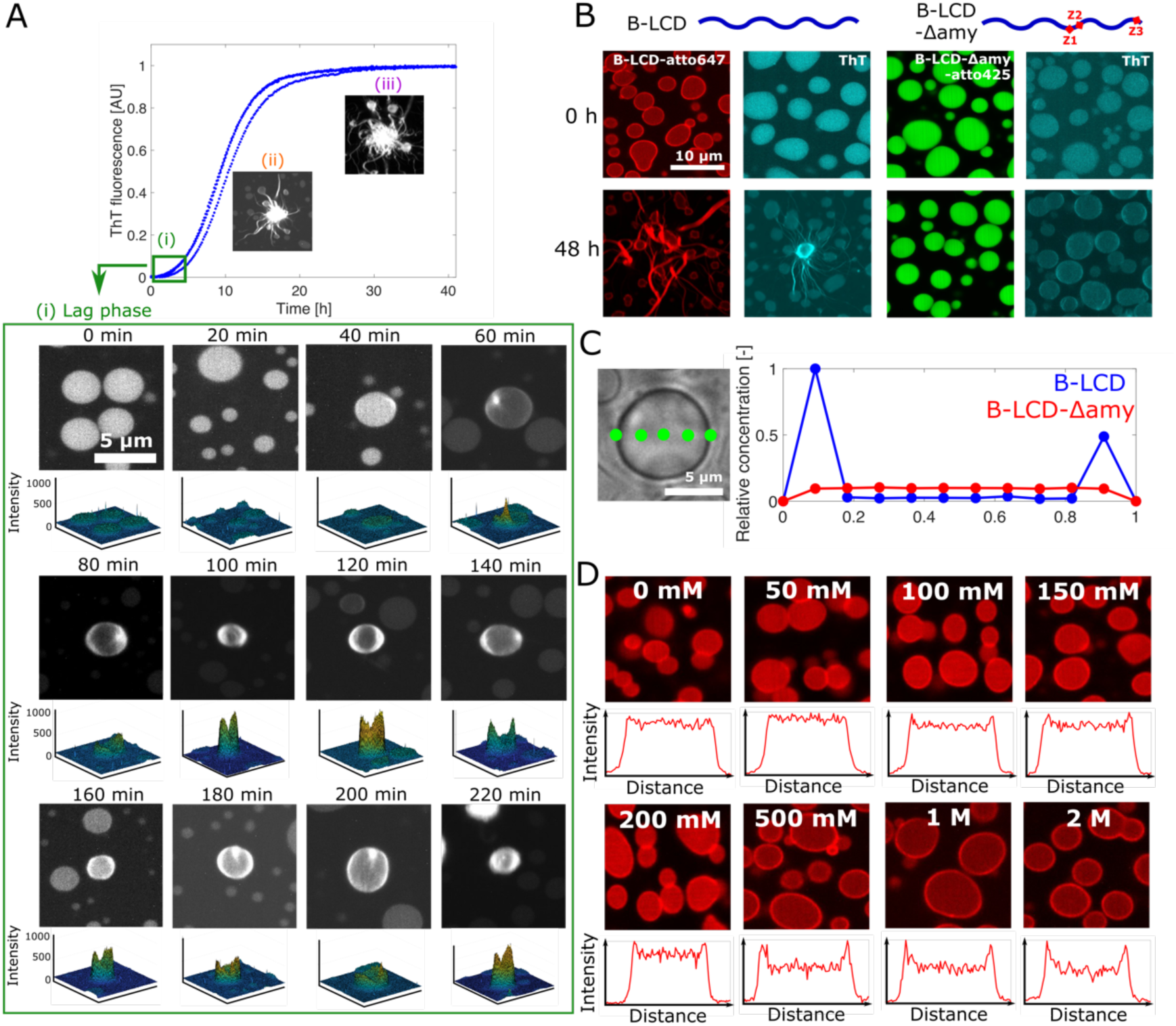
Formation of a protein-rich rim at the condensate interface precedes amyloid formation. **(A)** Re-scan confocal fluorescence microscopy images collected along the Thioflavin T profile. During the initial lag-phase (0-40 min) we observed the formation of the droplets and the uptake of ThT in their interior (i). Over time (60-220 min), a ThT-positive rim appeared on the droplet surface. 3D intensity profiles are depicted below each image, confirming higher ThT intensity value at the droplet edge. During the growth phase, star-shaped aggregates were formed from the droplets (ii, iii). Micrographs in (i-iii) are representative images taken at each time point. **(B)** Re-scan confocal fluorescence microscopy images of B-LCD labeled with atto647 at time 0 and after 48 h incubation. The rim at time 0 is not visible by ThT staining while after 48 h the high ThT signal at the interface indicates the presence of β-sheet-rich structures. In contrast, droplets formed by B-LCD-Δamy did not exhibit accumulation of molecules or fibrils at the droplet interface. **(C)** Protein concentration profile along the cross-section of an individual B-LCD droplet as measured by Raman spectroscopy, confirming the increase in protein concentration at the droplet surface. **(D)** Re-scan confocal fluorescence microscopy images of samples in (A) after 3 hours of incubation. The rim is more evident for samples at higher NaCl concentrations, which exhibit faster kinetics of amyloid formation (Fig. 2D). The intensity profiles show the normalized droplet intensity as a function of the normalized droplet edge-to-edge distance for one representative droplet.

To increase the resolution of the microscopy analysis, we labeled both B-LCD (with atto647) and B-LCD-Δamy (with atto425) and applied re-scan confocal fluorescence microscopy. Images of B-LCD samples taken a few minutes after preparation showed already the presence of a fluorescent rim along the droplet surface, which was not yet stainable by ThT (Fig. 4B). During incubation, the signal along this rim of individual droplets became ThT-positive, indicating the formation of fibrillar structures (Fig. 4B). Importantly, all droplets which eventually formed star-burst structures exhibited a ThT-positive rim, indicating that fibril formation at the condensate rim is an initial event during the liquid-amyloid transition of hnRNPA1-B LCD condensates (Fig. 4B).

To ensure that the formation of this rim was not an artifact related to labeling the B-LCD with a fluorophore, we measured the concentration profile along an individual unlabeled B-LCD droplet by Raman spectroscopy 30 min after droplet formation. The results confirmed the increase of protein concentration at the surface of the droplet formed by B-LCD but not by B-LCD-Δamy (Fig. 4C), consistent with the results of re-scan confocal fluorescence microscopy (Fig. 4B).

The rate of the liquid-amyloid transition increased with increasing salt concentration (Fig 2D). Consistently, re-scan confocal microscopy images acquired after 3 h incubation showed a more prominent rim for samples at higher salt concentrations (Fig. 4D).

In contrast to all the orthogonal data acquired with B-LCD, neither the increase in protein concentration at the droplet surface, nor the formation of this ThT-positive rim could be observed in the B-LCD-Δamy samples. Only a very weak ThT signal along the droplet surface was visible after 48 h incubation (Fig. 4B), consistent with the flat ThT kinetic profile acquired in bulk (Fig. 2A).

Overall, these results demonstrate that the formation of amyloid fibrils in B-LCD droplets is promoted at the surface of the condensates. Moreover, these important findings indicate the surface as a potential target to arrest the transition from liquid droplets into amyloids.

To confirm this hypothesis, as proof of concept, we aimed at delaying fibril formation by targeting the surface with two complementary strategies. In a first approach, we supplemented protein samples with 0.03% sodium dodecyl sulfate (SDS), one of the most common tensioactive agents. We monitored amyloid formation by ThT staining and fluorescence microscopy. The surfactant did not affect LLPS and the formation of liquid droplets, but drastically inhibited the increase of ThT signal and the formation of star-shaped aggregates over time (Fig. 5A-C).

**Figure 5:**
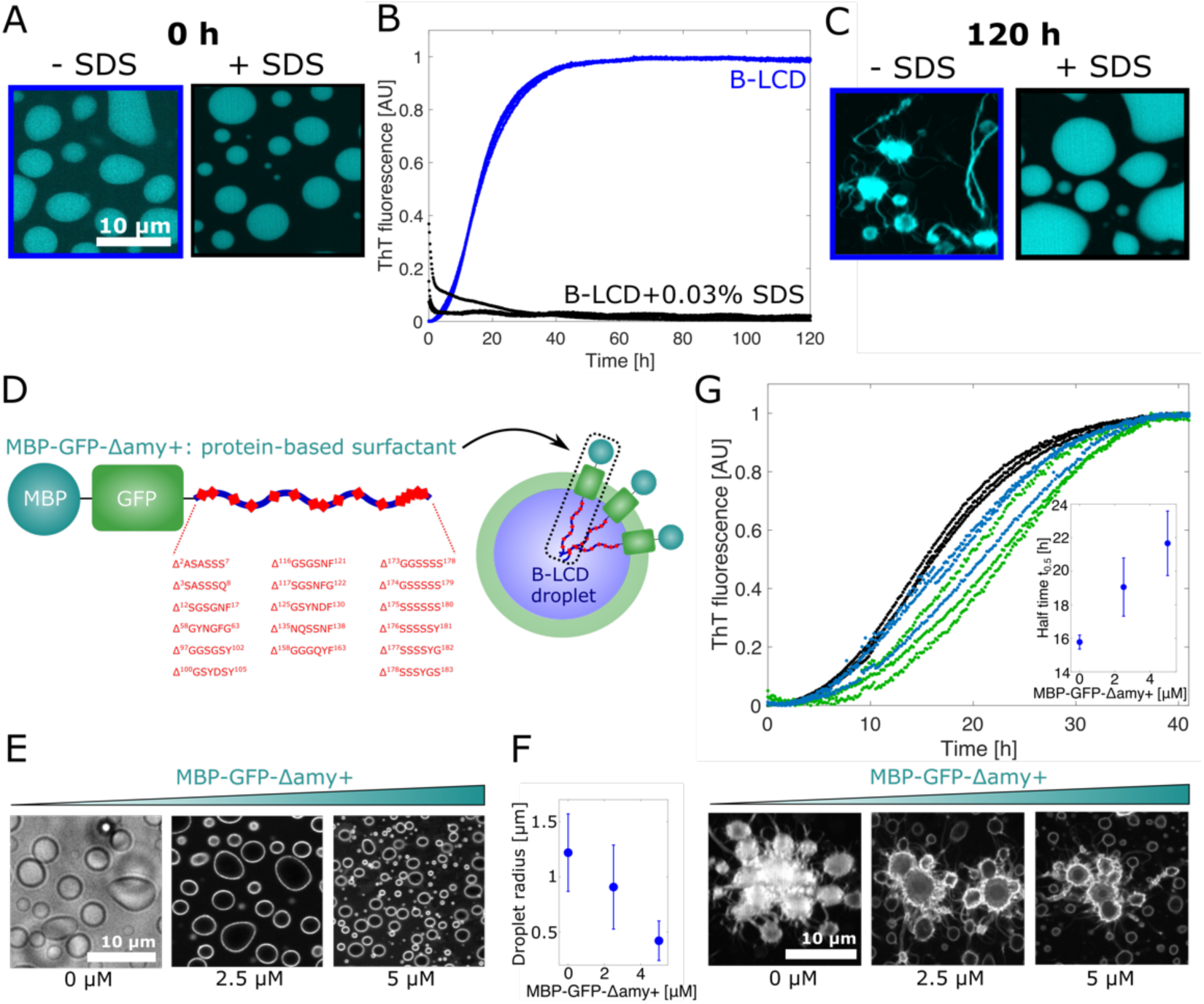
Targeting the surface of condensates inhibits amyloid formation. (**A-C**) The addition of 0.03% SDS to a 30 µM B-LCD solution did not affect LLPS (A) but prevented the increase of ThT signal over time (B) as well as the formation of rims and star-shaped aggregates after 120 h, as shown by re-scan confocal microscopy images (C). **(D)** Design of a protein-based surfactant (MBP-GFP-Δamy+) consisting of soluble Maltose-binding protein (MBP), Green-fluorescent protein (GFP) and a B-LCD variant lacking all predicted steric zippers (Δamy+). **(E)** Re-scan confocal fluorescence microscopy images showing accumulation of the protein-based surfactant molecules at the droplet surface. The fluorescent signal is originating from the GFP domain of the protein surfactant. Images were acquired at time 0. **(F)** Decrease of the average droplet size with increasing MBP-GFP-Δamy+ concentration. Error bars represent the standard deviation of sizes of > 1000 droplets from 3 independent samples. **(G)** In addition to changing the size of the condensates, increasing concentrations of the protein-based surfactant delays amyloid formation inside condensates. The inset depicts the average half times t_0.5_ extracted from the ThT profiles as a function of the MBP-GFP-Δamy+ protein surfactant. Black – 0 µM MBP-GFP-Δamy+, blue – 2.5 µM MBP-GFP-Δamy+, green – 5 µM MBP-GFP-Δamy+. Error bars represent the standard deviation of three independent samples. Re-scan confocal fluorescence microscopy images of samples after 40 h of incubation with increasing protein surfactant concentration.

In a second approach, inspired by the strategy described by Kelley et al.^54^, we generated a protein surfactant consisting of multiple domains: two soluble domains (Maltose-binding protein (MBP) and Green-fluorescent protein (GFP)) coupled to a B-LCD variant that lacks many segments predicted to form amyloids (indicated in the following as Δamy+) (Fig. 5D, Suppl. Table S2). This protein is expected to accumulate at the interface, with the Δamy+ LCD facing the interior of the condensates and the soluble domains exposed to the solution (Fig. 5D). Re-scan confocal microscopy analysis confirmed preferential accumulation of the surfactant protein at the interface of the droplets (Fig. 5E) and the decrease of the average size of the droplets in the presence of increasing concentrations of the protein surfactant (Fig. 5F). Fibril formation inside condensates coated with this protein was delayed, as demonstrated by the increase of the half times t_0.5_ (Fig. 5G), confirming that interfaces are crucial for fibril formation from condensates of the LCD of hnRNPA1.

## Discussion and conclusions

The LCD of hnRNPA1 exhibits a peculiar “molecular grammar”, comprising regions with sticker-spacer architecture that promote liquid-liquid phase separation^41,42^ and segments (LARKS) that induce fibril formation. Amyloid formation and liquid-liquid phase separation (LLPS) appear to be two independent processes, since disease-causing mutants show LLPS behavior similar to wild-type hnRNPA1^13^. Moreover, fibrillization is not required for LLPS^13^. We note that in general this can be different for other types of phase separation mechanisms, such as the one observed in liquid-liquid crystalline phase separation (LLCPS)^55^.

Here we analyzed the C-terminal LCD of two splicing variants of hnRNPA1, isoforms hnRNPA1-A (A-LCD) and hnRNPA1-B (B-LCD). As control, we generated a sequence lacking three important segments promoting amyloids. We observed that wild-type and mutant LCD have essentially identical LLPS behavior (Fig. 1). However, amyloid formation is drastically reduced for the mutant LCD lacking three fibril forming segments (Fig. 2). Therefore, our data demonstrate that liquid-liquid phase transition and fibrillization are connected but distinct processes which are independently mediated by different regions of the protein sequence. This is also consistent with our finding that LLPS is insensitive to changes in the ionic strength, while fibril formation is accelerated at increasing salt concentrations. These observations demonstrate that the two processes are governed by different interactions which can be independently modulated in different ways (Fig. 1E and 2E).

Even though LLPS and fibrillization are decoupled processes, LLPS strongly promotes the latter (Fig. 3). Indeed, at concentrations lower than the LLPS saturation concentration (2.5 µM), fibril formation was not observed within seven days. Condensation is therefore required for fibrillation or at least strongly accelerates fibril formation. A first intuitive explanation of this observation is the local increase of protein concentration of approximately 1000-fold inside the condensates. This high concentration can have both a thermodynamic and a kinetic effect. In test tubes, proteins containing steric zippers have typical critical concentrations for amyloid formation in the nano- and micromolar range^56^, well below the concentration inside the condensates (in the millimolar range) (Fig. 1E). From a kinetic point of view, the local increase in protein concentration can accelerate amyloid formation by increasing nucleation and growth rates^57,58^.

However, our analysis of the spatial evolution of fibrillation inside the condensates shows that fibril formation is promoted at the interface of the condensates (Fig. 4), indicating that the interplay between condensation and fibril formation is beyond a simple increase of local protein concentration.

Previous studies have shown that proteins can exhibit different conformations inside and outside the condensates. For instance, the protein tau adopts a more extended conformational ensemble within droplets which exposes an aggregation-prone region, therefore promoting amyloid formation^22^.

In addition to conformational changes inside and outside the condensate, Farag et al. have recently shown that LCD molecules of hnRNPA1 are organized in different topologies inside the condensates^59^. Specifically, proteins are more expanded at the interface compared to both the interior of the condensate and the dilute phase^59^. Moreover, molecules prefer to be oriented perpendicularly to the condensate interface^59^.

These observations provide a plausible explanation of the promotion of amyloid formation at the interface of the condensates reported in this work. The orientation and conformation expansion of the molecules at the interface likely promotes β-sheet formation of aggregation-prone regions of the hnRNPA1 proteins which are brought in close proximity at the interface.

This process shares common aspects with surface-induced protein aggregation in homogeneous solutions, where interfaces are generally well-known to promote heterogeneous nucleation events by inducing protein adsorption, local increase of protein concentration and possible conformational changes which trigger aggregation^60–65^.

In addition to promoting conformational changes and orientation^59^, interfaces of condensates can preferentially partition components^66^ and accelerate or inhibit fibrillation of client molecules^67^. Overall, interfaces can therefore represent a special location for biochemical activity within condensates and contribute to the complex interplay between condensation and fibrillation. Local increase of concentration increases polymerization and aggregation rate in the condensates, as observed for instance with actin^68^. However, condensates are not simple liquids but viscoelastic materials^4^ composed of multiple components. Heterotypic buffering^39^ can prevent amyloid formation of both scaffold^39^ and client proteins^69^ in the condensates despite the high local concentration. Interactions and conformations can differ at the interface with respect to the bulk of the condensate, further modulating the interplay between condensation and amyloid formation.

Consistent with our findings that fibrillation of the LCD of hnRNPA1 is promoted at the interface of condensates, we have demonstrated that the manipulation of the condensate interface affects the kinetics of fibril formation, resulting in a delay or complete arrest of amyloid generation (Fig. 5).

The results that we have presented here (Fig. 6) could have implications for the development of therapeutic strategies aimed at arresting aberrant fibril formation mediated by condensation and indicate condensate interfaces as a potential therapeutic target.

**Figure 6:**
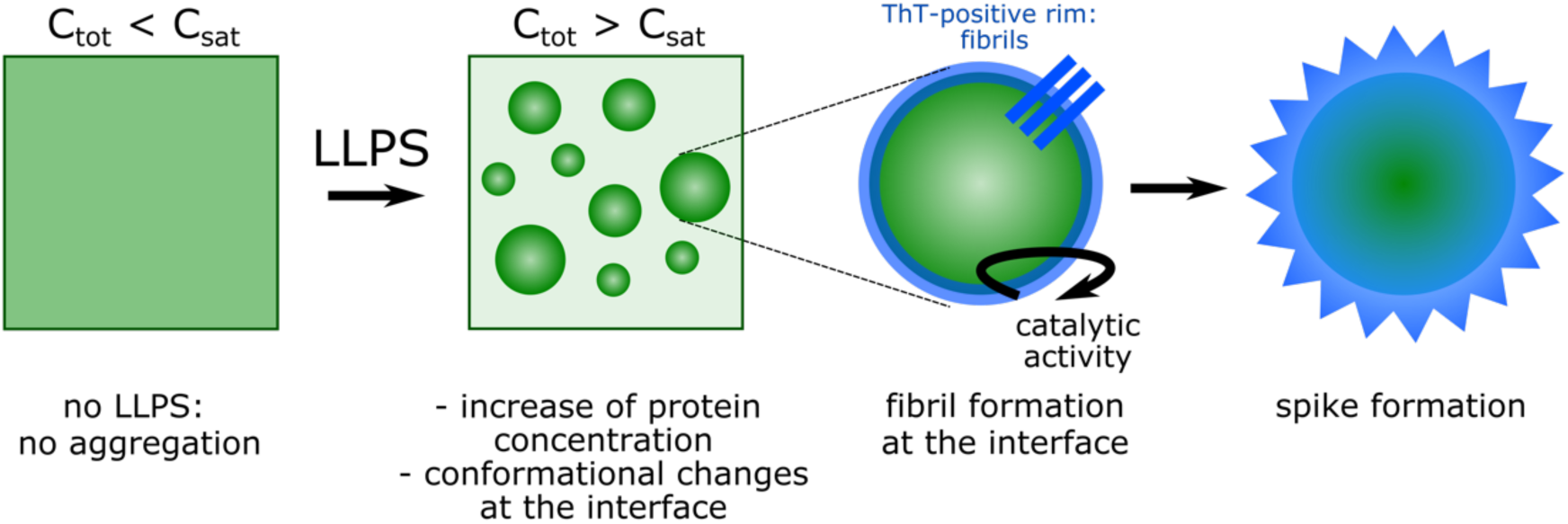
Schematic illustration of amyloid formation promoted at the interface of condensates of the LCD of hnRNAP1.

## Acknowledgements

The authors gratefully acknowledge financial support from the Swiss National Science Foundation (grants 205321_179055), the Synapsis Foundation and the Claude and Giuliana Foundation. We gratefully acknowledge Rohit Pappu and Mina Farag for helpful discussions.

## Author contributions

ML, LF, UCP and PA contributed to the conceptualization of the work, ML, LF, UCP, CS, AMK, DP and JZ performed the experiments, ML, JZ, RM and PA analyzed the data, ML and PA wrote the initial manuscript, everybody reviewed and edited the manuscript, PA supervised the work and acquired funding.

## Competing interests

The authors declare no competing interests

## Materials and Methods

### Protein expression and purification

hnRNPA1-A-LCD (13.1 kDa, sequence shown in Fig. 1A), hnRNPA1-B-LCD (17.6 kDa, sequence shown in Fig. 1A), hnRNPA1-B-LCD-Δamy (15.8 kDa, sequence shown in Fig. 1A) and MBP-GFP-Δamy+ were recombinantly expressed and purified in *Escherichia coli* BL21 DE3 cells. All DNA sequences were synthesized, codon optimized for expression in E. coli, and cloned into the pET-15b vector by Genewiz (NJ, US). Plasmids containing the sequences for the different constructs, including N-terminal 6x-His tags, and IPTG-inducible promotor as well as ampicillin resistance, were transformed *via* heat shock at 42 °C for 30 s. Cells were grown at 37°C on LB agar plates containing 100 µg/ml ampicillin and further scaled up in rich media. For the MBP-GFP-Δamy+ surfactant, conventional Luria Broth media was used. Protein expression was induced by addition of 0.5 mM IPTG and incubation overnight. Following harvesting by centrifugation, the cells were re-suspended in lysis buffer (8 M urea, 1M NaCl, 50 mM Tris, pH 7.5), lysed by sonication, and centrifuged at 12 800 RPM for 20 min. The supernatant was transferred onto a Nickel NTA column, washed with washing buffer (1M NaCl, 50 mM Tris, pH 7.5, 50 mM imidazole) and eluted by increasing the imidazole concentration in the buffer to 500 mM. The eluate was further polished by size exclusion chromatography on a Superdex 75 column (Cytiva) using a buffer containing 8 M (for A-LCD, B-LCD-Δamy) or 2 M urea (for B-LCD), 1 M NaCl, 50 mM Tris, pH 7.5, 10 % glycerol and 2 mM β-mercaptoethanol. The eluted fractions were tested by SDS-PAGE and Coomassie blue staining and pure fractions were pooled and immediately used for analysis. A representative chromatogram and gel are shown in Suppl. Fig. S6. Typically, the LCD and LCD-Δamy proteins were concentrated to 200 µM and 600 µM stock solutions, respectively, using spin filters (Merck, Millipore, MWCO: 5 kDa).

The protein-based surfactant MBP-GFP-Δamy+ construct was expressed, harvested and purified in the same way but by using urea-free buffers containing a reduced NaCl concentration of 500 mM. Size exclusion chromatography was carried out on a Superdex 200 column (Cytiva).

For labeling experiments, LCD and LCD-Δamy were expressed and purified as described before, but the Tris-containing SEC buffer was replaced by phosphate-buffered saline supplied with 1 M NaCl and 2 M or 8 M urea. Following the instructions of the manufacturer, the purified protein samples were supplied with 2 mg/ml atto647 or atto425 dye dissolved in 100 % DMSO and incubated overnight at room temperature. Subsequently, the free dye was separated from the atto-conjugated proteins by size exclusion chromatography on a Superdex 75 column (Cytiva).

### Phase separation, fluorescence imaging and confocal microscopy

The phase separation of the B-LCD and the B-LCD-Δamy proteins was induced in 384-well plates (Matriplate, Brooks).

Brightfield and widefield fluorescence microscopy were carried out on a Nikon Ti2 Eclipse inverted microscope, equipped with an LEDHub light source (Omicron) and an Andor Zyla camera, using a 60x oil objective (Nikon, NA = 1.4).

Confocal microscopy was performed with a Nikon NSTORM (Nikon UK, Ltd) system equipped with a Rescan Confocal Microscope RCM1 (Confocal.nl, Amsterdam, the Netherlands). We used an sCMOS camera (Orca Flash 4.0 V2) and a Nikon SR Apochromat TIRF objective 100x / 1.49 with oil immersion. The different laser excitations were at 488 nm, and 647 nm, in order to excite the ThT, atto488 and atto647 dyes, respectively. The setup was fully controlled, and image acquisition was performed using the NIS-Elements software (Nikon). The implemented re-scan unit provides an enhancement in resolution from 240 nm to 170 nm^70,71^.

### Amyloid formation kinetics

Amyloid formation was monitored by recording the fluorescence intensity of Thioflavin T (ThT) over time. To this aim, samples were supplied with 20 µM ThT and measured in 384-well plates (Matriplate, Brooks) on a ClarioStar plus plate reader (BMG Labtech). To avoid evaporation, the plates were sealed with sticky aluminum foil (Corning). Measurements were taken every 10 minutes over several days, by applying excitation at 450 nm and measuring emission at 490 nm.

### Raman spectroscopy

Raman spectra were acquired on a confocal inverted microscope (Nikon Ti-E) connected to a Raman system (Horiba, LabRAM HR Evolution UV-VIS-NIR). The sample was excited using a 532 nm laser (Nd:YAG, Cobolt Samba™ cw single frequency) at a power of 75 mW and the Raman signal was detected with a Synapse EM-CCD detector (Horiba).

The protein concentration inside the droplets was measured as previously described^50^. In brief, a characteristic peak at a wavenumber of 1000 cm^-1^ corresponding to the symmetric breathing of the benzyl ring of phenylalanine was used as internal standard. The intensity of this peak corresponds to the number of phenylalanines in the sample, which can be directly related to the protein concentration by a standard curve.

### Phase separation and liquid-solid transition in microfluidic water-in-oil droplets

Fabrication of master wafers and polydimethylsiloxane (PDMS) devices were performed by standard soft lithography techniques as previously described^48^. The resulting devices exhibited a flow focusing architecture containing three inlets, two mixing junctions and one outlet. At the first junction, a solution of homogeneous B-LCD (120 µM stock) in 50 mM Tris, pH 8.5, 2M urea, 2 mM βmercaptoethanol was mixed with a buffer containing 50 mM Tris at pH 7.5 to induce phase separation. At the second junction, this mixture was encapsulated into water-in-oil droplets using an HFE-7500 oil (3M) supplied with 0.5% Pico-surf surfactant (Sphere Fluidics). To observe the droplets over hours or days, the water-in-oil compartments containing the phase separated droplets were transferred into glass capillaries and tightly sealed. Images were taken by brightfield or epi-fluorescence microscopy.

### Atomic Force Microscopy (AFM)

AFM imaging was carried out by a Nanoscope Multimode 8 scanning probe microscope (Bruker, USA). 50 µl of a freshly prepared B-LCD droplet solution was deposited and then adsorbed onto freshly cleaved mica for 5 min before imaging at room temperature. The imaging was operated under hydrated condition in a liquid cell with a V-shaped silicon nitride cantilever (Bruker, USA) with a nominal spring constant of 0.7 N/m and tip radius of 2 nm.

The AFM probe was scanned at a scan frequency of 0.4 Hz on the sample continuously to screen the evolution of microdroplets. AFM images were flattened using the Nanoscope 8.1 software (Bruker, USA), and no further image processing was applied.

### Zipper and LARKS prediction using ZipperDB and LARKSdb

Steric zipper regions were predicted using the data base ZipperDB^72^ available online at https://services.mbi.ucla.edu/zipperdb/.

LARKS were predicted using the LARKSdb^43^ available online at https://srv.mbi.ucla.edu/LARKSdb/index.py.

## Notes

### Competing Interest Statement

The authors have declared no competing interest.

