## Supplementary Information for "The interface of condensates of the hnRNPA1 low complexity domain promotes formation of amyloid fibrils"

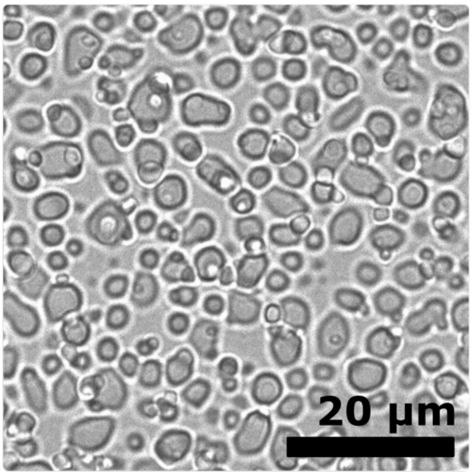


**Supplementary Figure 1:** The LCD of the isoform A of hnRNPA1 (A-LCD) undergoes liquid-liquid phase separation.

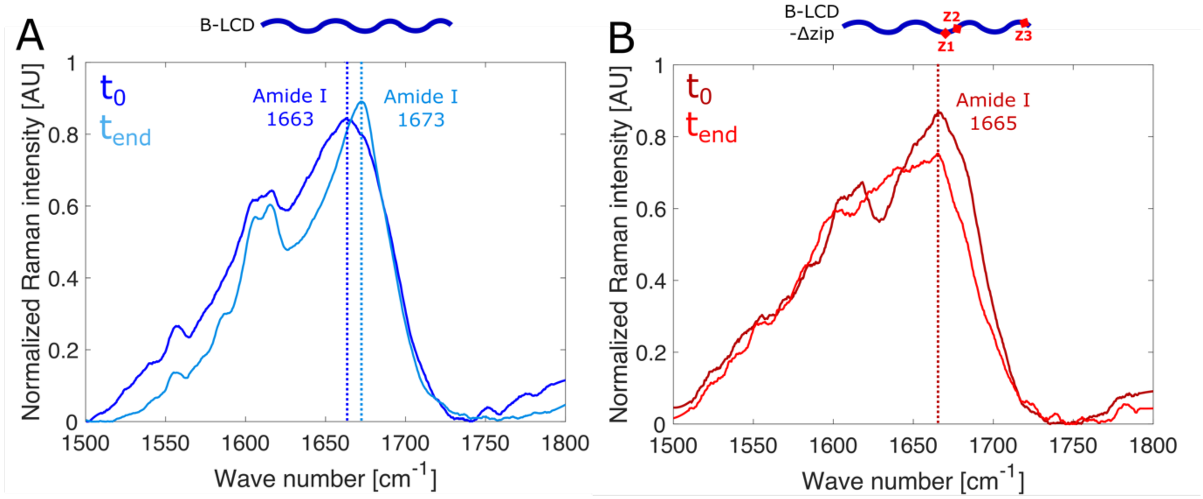


**Supplementary Figure 2:** Raman spectra (Amide I regions) of B-LCD (left panel) and B-LCD-Δamy (right panel) before and after incubation, showing variation in the Amide I region. B-LCD exhibited a shift of the Amide I peak from lower to higher wave numbers (1663 to 1673 cm$^{-1}$) which was absent in B-LCD-Δamy.

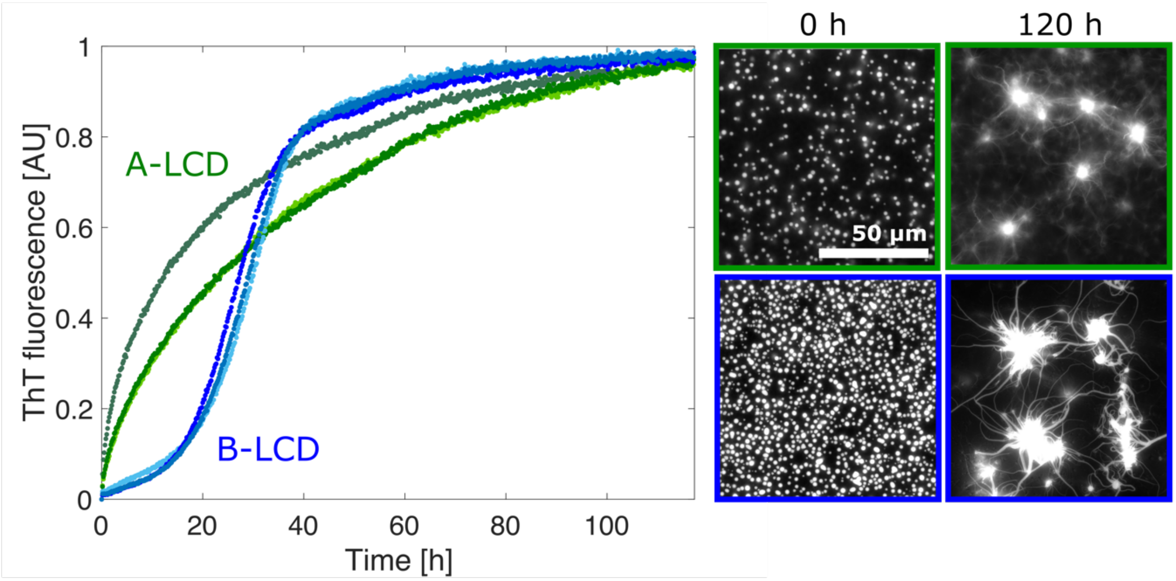


**Supplementary Figure 3:** Comparison of the liquid-amyloid transition of A-LCD and B-LCD. Both proteins undergo liquid-amyloid transition and form starburst structures over time. However, A-LCD aggregates faster than B-LCD.

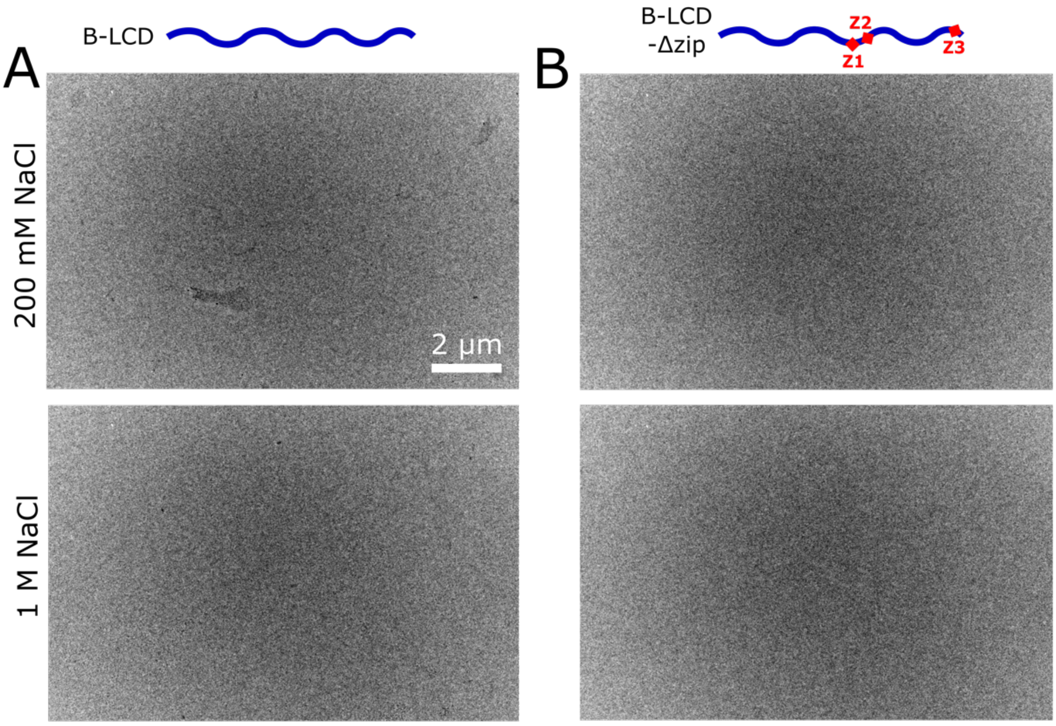


**Supplementary Figure 4:** TEM images after incubation for 4 days at sub-critical LLPS concentrations. Neither the B-LCD nor the B-LCD-Δamy construct shows higher order structures.

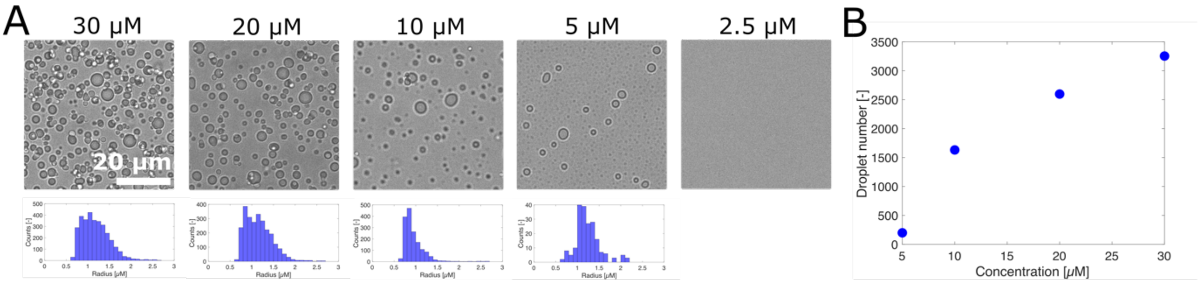


**Supplementary Figure 5: (A)** Images of of B-LCD droplets formed at various initial protein concentrations acquired by brightfield optical microscopy and corresponding size distributions extracted from these images. **(B)** Number of droplets extracted from images in (A).

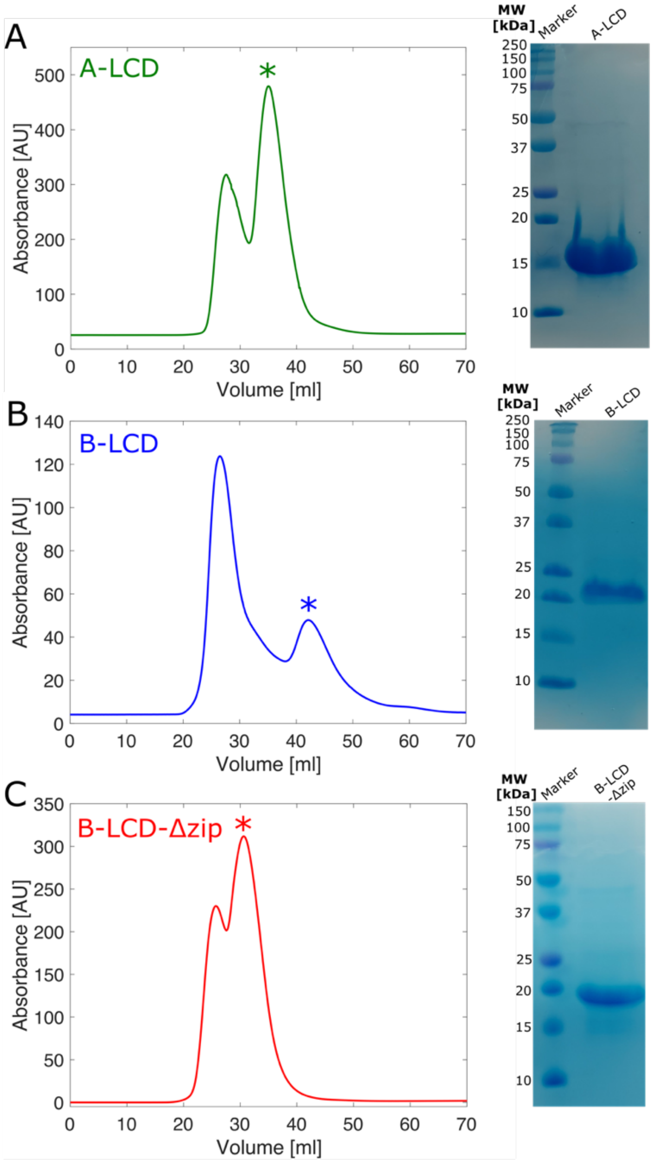


**Supplementary Figure 6:** Purification of A-LCD (A), B-LCD (B) and B-LCD-Δamy (C) by size exclusion chromatography. Asterisks in chromatograms show fractions considered for the experiments. Purity was verified by poly-acrylamide SDS-PAGE, followed by Coomassie staining. All constructs show the expected molecular weight and high purity.

**Supplementary Table 1:** Steric zippers of the LCD of hnRNPA1-B, as predicted by ZipperDB. Highlighted in yellow are steric zipper sequences which have been deleted to obtain the less-aggregation prone variant B-LCD-Δamy. Steric zipper regions which overlap with LARKS are marked with an asterisk (*).

| Position | Sequence | Rosetta Energy | Shape Compl. | Area of Interface | Contact Area | SASA | C-Score |
|---|---|---|---|---|---|---|---|
| 176 | SSSSSY* | -26.2 | 0.82 | 70 | 0 | 191 | -45.45 |
| 3 | SASSSQ* | -24.8 | 0.82 | 70 | 0 | 194 | -44.05 |
| 175 | SSSSSS* | -24.6 | 0.82 | 70 | 0 | 192 | -43.85 |
| 177 | SSSSYG* | -24.6 | 0.841 | 79 | 0 | 204 | -45.07 |
| 2 | ASASSS* | -24.4 | 0.802 | 72 | 0 | 194 | -43.58 |
| 174 | GSSSSS* | -24.4 | 0 | 0 | 0 | 187 | 0 |
| 125 | GSYNDF | -24.2 | 0.833 | 61 | 0 | 177 | -42.75 |
| 116 | GSGSNF* | -24 | 0.894 | 35 | 0 | 176 | -40.86 |
| 117 | SGSNFG* | -24 | 0.792 | 78 | 0 | 196 | -43.63 |
| 58 | GYNGFG* | -23.8 | 0.833 | 67 | 0 | 200 | -43.00 |
| 135 | NQSSNF | -23.6 | 0.732 | 90 | 0 | 208 | -43.53 |
| 178 | SSSYGS* | -23.6 | 0.796 | 60 | 0 | 187 | -41.54 |
| 100 | GSYDSY* | -23.4 | 0.835 | 56 | 0 | 177 | -41.48 |
| 158 | GGGQYF | -23.4 | 0.848 | 64 | 0 | 199 | -42.52 |
| 173 | GGSSSS* | -23.3 | 0.701 | 58 | 0 | 182 | -39.57 |
| 97 | GGSGSY* | -23.2 | 0.701 | 58 | 0 | 185 | -39.47 |
| 12 | SGSGNF* | -23.2 | 0.736 | 78 | 0 | 194 | -42.04 |

**Supplementary Table 2:** Sequence of the Δamy+ LCD, lacking all predicted steric zipper regions (marked in yellow).

| | |
|---|---|
| **B-LCD** | 1 11 21 31 41 51<br>MASASSSQRG RSGSGNFGGG RGGGFGGNDN FGRGGNFSGR GGFGGSRGGG GYGGSGDGYN<br>61 71 81 91 101 111<br>GFGNDGGYGG GGPGYSGGSR GYGSGGQGYG NQGSGYGGSG SYDSYNNGGG GGFGGGSGSN<br>121 131 141 151 161 171<br>FGGGGSYNDF GNYNNQSSNF GPMKGGNFGG RSSGPYGGGG QYFAKPRNQG GYGGSSSSSS<br>181<br>YGSGRRF |
| **B-LCD-Δamy+** | 1 11 21 31 41 51<br>MRGRGGGRGG GFGGNDNFGR GGNFSGRGGF GGSRGGGGYG GSGDNDGGYG GGGPGYSGGS<br>61 71 81 91 101 111<br>RGYGSGGQGY GNQGSGYNNG GGGGFGGGGG NYNGPMKGGN FGGRSSGPYG AKPRNQGGYG<br>121<br>RRF |
